# Learning structure in gene expression data using deep architectures, with an application to gene clustering

**DOI:** 10.1101/031906

**Authors:** Aman Gupta, Haohan Wang, Madhavi Ganapathiraju

## Abstract

Genes play a central role in all biological processes. DNA microarray technology has made it possible to study the expression behavior of thousands of genes in one go. Often, gene expression data is used to generate features for supervised and unsupervised learning tasks. At the same time, advances in the field of deep learning have made available a plethora of architectures. In this paper, we use deep architectures pre-trained in an unsupervised manner using denoising autoencoders as a preprocessing step for a popular unsupervised learning task. Denoising autoencoders (DA) can be used to learn a compact representation of input, and have been used to generate features for further supervised learning tasks. We propose that our deep architectures can be treated as empirical versions of Deep Belief Networks (DBNs). We use our deep architectures to regenerate gene expression time series data for two different data sets. We test our hypothesis on two popular datasets for the unsupervised learning task of clustering and find promising improvements in performance.

## Introduction

Genes and proteins play a crucial role in the physiology of all cellular organisms. Thus, studying the behavior of genes and their products under the influence of stimuli is an important goal in biology. Genes influence each other via their products like messenger RNA (mRNA) and proteins. Measuring the amount of mRNA of genes in a system is a useful way to quantitatively study the interdependence in a set of genes. Fortunately, DNA microarray technology has changed the field of genomics by providing a cost-effective and fast mechanism to measure the mRNA expression levels of thousands of genes in a single experiment. This gene expression data has proved useful for a number of supervised and unsupervised machine learning tasks. For example, gene regulatory network inference is an open and challenging problem that exploits gene expression data. Several methods have been proposed to infer gene regulatory network inference [1][2][3]. Another example is the clustering of gene expression data to group similar genes together for better understanding of interactions [4].

Gene expression data is consumed in the form of either time-series expression of a number of genes, gene expression profiles of different patients/organisms or the steady-state expression values of genes under varying degrees of perturbation. In the case of steady-state data, each perturbation experiment and resulting gene expression vector is equivalent to the gene expression profile of an individual organism. In this paper, we propose using multiple samples to capture interesting characteristics of the underlying distribution of gene expression vectors. For example, the interaction between gene A and B might be exhibited via high correlation in gene expression values. To facilitate the task of learning interesting properties of the input distribution, we turn to deep learning, a field that has taken big strides over the last few years. Recent advances have made it possible to use deep architectures to facilitate supervised and unsupervised learning tasks. We explore using deep architectures, trained using denoising autoencoders, for the task of learning a low-dimensional representation of gene expression profiles. Generative models like Deep Belief Networks (DBNs) are well suited for this task [5][6], but are difficult to train owing to difficulties in computing partition functions. We propose stacking together layers of a special variant of autoencoders, referred to as denoising autoencoders (DAs) [7], to initialize a deep neural network. Denoising autoencoders have proved useful for extracting features from data by making the learned representation resistant to partial corruption of the input. In the process of training, the denoising autoencoder learns to “guess” missing or corrupted values. The stacked version of this architecture is also referred to as stacked denoising autoencoder (SDA) [8].

Once the network has been trained, we treat the lowdimensional representation of latent input examples as a collection. Since denoising autoencoders have proved successful in learning features for tasks like image classification [8], we claim that learned deep architectures could generalize interesting characteristics of the underlying input distributions. To test the hypothesis that the network has learnt interesting characteristics of this distribution, we use the collection of low-dimensional representations as an empirical distribution and regenerate gene expression profiles. Since gene clustering has important applications, we then test the above-mentioned hypothesis by comparing raw input data and regenerated data on the task of clustering. The performances of both data on the task of clustering are compared with the help of gold standard cluster labels for the genes under consideration.

## Background

### Related work and motivation

Microarray data denoising and enhancement has received considerable attention from the research community over the past several years. An example is the usage of wavelet transforms to enhance microarray images [11]. Missing value imputation is another area of interest. Successful attempts include using a Bayesian method for imputation of missing values [9] and using a least squares based method [10].

Another initial work is [15], they applied the standard PCA on gene expression data and studied the effectiveness of principal components in capturing cluster structure. Their results showed that knowledge captured by principal components does not necessarily improve cluster quality. In a recent work [20], denoising autoencoder is applied to effectively summarize key features in breast cancer data.

In this work, we do not aim at removing noise from microarray data. Rather, our objective is to learn interesting patterns in the input distribution of gene expression profiles. We aim to learn these interesting characteristics and generalize them across all training samples. We also aim at demonstrating the efficacy of using this enhanced data for a popular unsupervised learning task involving clustering genes into groups. For the task of enhancing gene expression data, we use deep networks trained via denoising autoencoders.

We now describe the generic autoencoder device so as to acquaint the reader about the specific properties of the deep network.

### The Autoencoder

Autoencoders are a class of neural networks that attempt to learn a compact representation of data. Assume that an input gene expression profile is represented by ***x*** ∈ [**0**, **1**]***^d^***, drawn from a distribution ***P*** of expression profiles. The autoencoder network first maps the ***d*** – dimensional input ***x*** to a representation 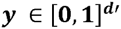 via the following method:

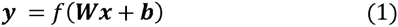

Here, *f* is a non-linear function like the sigmoid or tanh (hyperbolic tan). ***W*** is a ***d***′ ***x d*** dimensional matrix and ***b*** is a bias term. once this mapping has been performed, the embedding is mapped back into an output **z** of **d**-dimensions, the same shape as that of **x**. The mapping is done in the following manner:

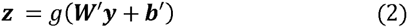

Here, ***g*** is again a non-linearity like sigmoid or tanh. ***W****′* is a ***d x d′*** dimensional matrix. More often than not, both sets of weights are constrained such that ***W***′ **= *W***. ***z*** is a reconstruction of the input ***x***, by first transforming to ***y*** and then expanding/contracting back. The network is then trained to minimize the average reconstruction error. The parameters of this model are **W**, **b** and **b**'; and if one doesn’t use tied weights, also **W**'.

Note that it is not necessary that ***d'*** < *d*. In general, however, the hidden layer does not have higher dimensions because of the problem of overfitting, unless regularization is used. If ***d*′ = *d***, then the network can learn the identity mapping and the latent representation generated will not be very interesting. When the hidden layer is more compact than the input layer and the function *f* is linear, the hidden representation is equivalent to performing Principal Components Analysis (PCA) on the input [13]. Things start becoming interesting when ***f*** is non-linear. In that case, the network has a chance to minimize the reconstruction error by opting to go into a non-linear space. Thus, the hidden representation becomes different from PCA and can be viewed as a low-dimensional coding of the input representation learned via non-linear transformations.

The reconstruction of input can be cast as an optimization problem. Among several popular loss functions, squared error is popular:

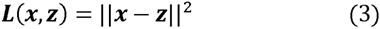

Another popular loss function is the cross-entropy of the reconstruction:

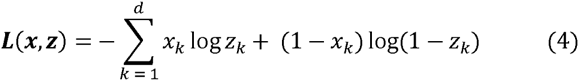

Standard optimization algorithms like stochastic gradient descent (SGD) can be used to optimize the loss function.

Autoencoders have been successfully used as initialization blocks of deep networks [12] and for dimensionality reduction [14].

## Methods

In this section, we propose using stacked denoising autoencoders to initialize deep architectures. First, we present the denoising autoencoder and its unique characteristics including types of training noises. Then we formalize the deep architecture used to learn low-dimensional encodings of the input training examples. Subsequently, we describe generation of samples from the architecture akin to generating samples from architectures like deep belief networks. We describe the deep architecture used to learn low-dimensional codes of gene expression profiles. Finally, we assess the quality of these samples on the task of clustering on gold standard labels.

### A. Using the Denoising Autoencoder (DA) for gene expression data

Autoencoders are trained to reconstruct input data using an intermediate representation. our objective is to extract multimodal relationships and complex patterns in the distribution of gene expression profiles without requiring explicit knowledge about the domain.

One strategy to force the network not to learn an identity mapping is to constrain the hidden layer such that **d’ < d**. This requires the network to learn a lossy compression of the input. Another novel strategy to modify plain vanilla autoencoders is to train the network to reconstruct partially corrupted input. In this process, the network is forced to “guess” missing/corrupted values. Autoencoders using this strategy are referred to as Denoising Autoencoders [7]. Specifically, input ***x*** is first corrupted (using a suitable corruption scheme) to generate 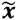. An intermediate representation ***y*** can be generated using the deterministic mapping 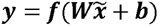, where ***W*** and ***b*** are parameters of the mapping. As described in the previous section, we can then reconstruct ***z* = *g*** (***Wy* + *b***′). Unlike the strategy used in the previous section, the reconstruction ***z*** uses 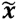 instead of ***x***. The training criterion used is still the same:

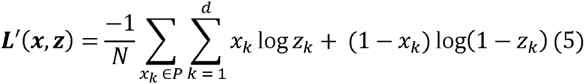

***L***′ is the function that minimizes the empirical risk over ***N*** samples drawn from the distribution ***P***. The network is still guided towards the clean signal ***x***. But to for every step taken towards lowering the cost of reconstruction, the network must learn weights that help it fill-in missing or corrupted values in the modified input.

The use of denoising autoencoders has met with considerable success, since the intermediate representation corresponds to useful and interesting features. Gene expression profiles are generally available as expression values for a set of genes. Learning the regulatory relationships between genes is an important goal in systems biology, and expression profiles have been extensively used for various inference algorithms. However, analysis of domains like image data via deep learning uses the spatial linkages of pixels. Generally, gene expression data does not have such relationships. But we argue that since the interdependencies between genes can be inferred from expression values, gene expression profiles lend themselves perfectly towards the task of estimating statistical relationships in the empirical distribution ***P***.

### Types of noise

Most methods used to enhance or denoise gene expression data make some assumptions or use domain knowledge to achieve satisfactory results. In this work, we make no such assumptions and do not utilize any biological knowledge. We consider some generic methods of corrupting input for training. Among these, additive Gaussian noise 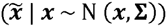 and masking noise are popular. Masking noise involves setting a fixed fraction of the bits of the vector ***x*** to zero. In this work, we focus on using masking noise to corrupt the input. The fraction of noise is controlled by the hyper-parameter *v*.

### The Denoising Autoencoder vs PCA

As discussed in the previous section, denoising autoencoders using non-linear transformation functions are fundamentally different from performing Principal Components Analysis on the input samples. PCA captures the directions of maximum variance in the data in the linear space. [15] demonstrates that performing PCA on gene expression data does not necessarily improve quality of clustering. We claim that the non-linear capabilities of DAs make them capable of learning more complex relationships in the input distribution. A demonstration of the same can be found in [14] for the task of image reconstruction.

### A deep architecture using multiple denoising autoencoders parallels

Denoising autoencoders can be stacked together to form stacked denoising autoencoders. Once the i^th^ layer has been trained, its output is used to train the (i+1)^th^ layer. Subsequently the entire deep network can be fine-tuned using a global loss function. This technique of greedy layer-wise tuning followed by global fine-tuning has yielded good results than using random weights as a starting point for global optimization [16].

Fig. 1 represents a stacked denoising autoencoder. For an n-deep SDA, there are (2n-1) hidden layers. The network can be visualized as an encoding of input, followed by decoding. The middle layer has the lowest dimensions and is known as the bottleneck layer.

**Fig. 1.**
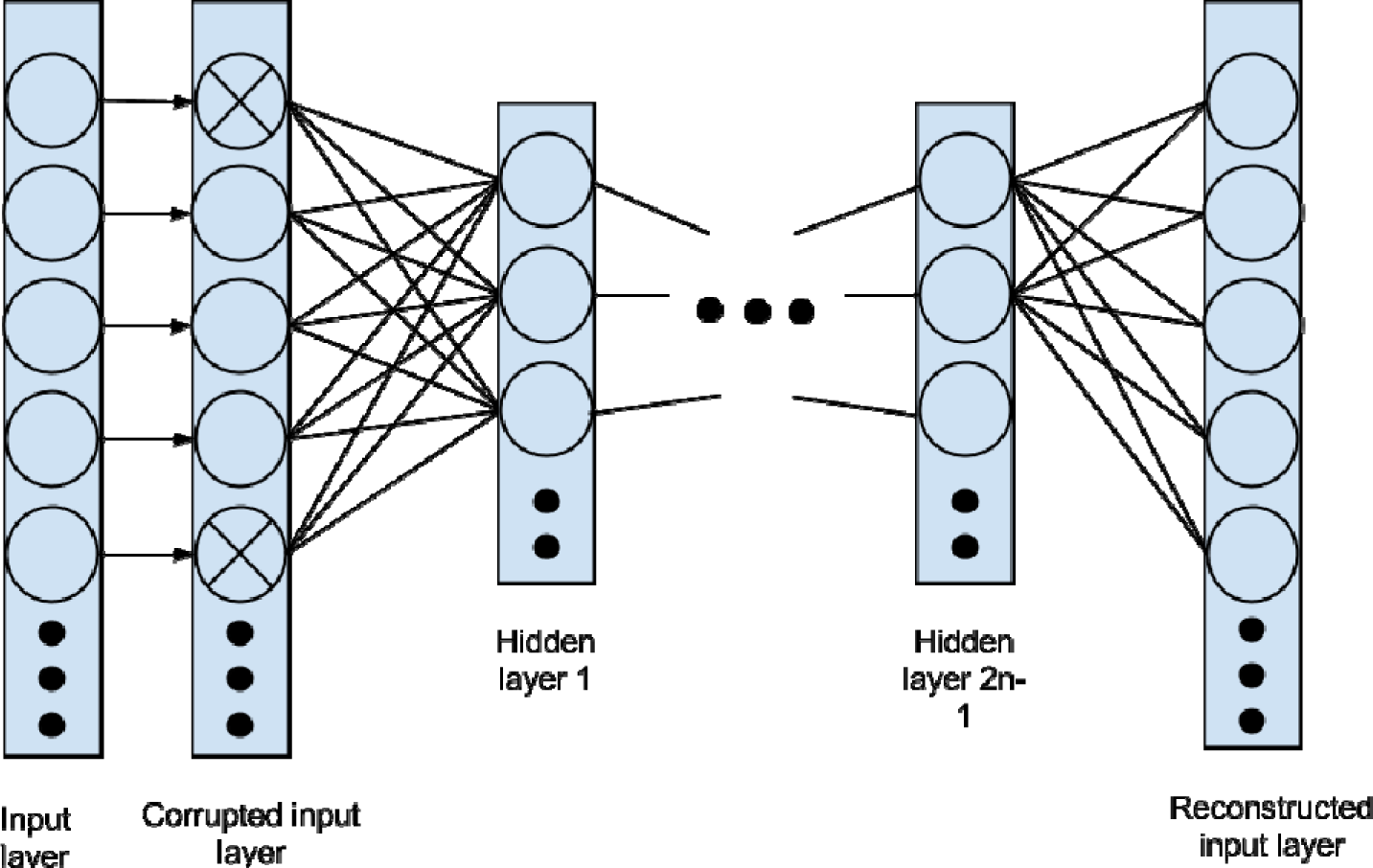
Depiction of a stacked denoising autoencoder. For an n-encoder, there are 2n-1 hidden layers. Individual layers can be pre-trained greedily before a final global fine-tuning is applied to the entire network.

### Drawing parallels to generative models

All transformations in an autoencoder-based network are deterministic in nature, and thus autoencoders are not generative models. However, they are similar to generative models like the Deep Belief Networks (DBNs). Thus, we can treat the higher-level representation of the training examples used to train the network as a generative empirical distribution. By successively decoding this representation using the decoding layers, we can generate samples from this empirical distribution. The regeneration of examples can help us compare the performance of the network to the performance of raw input data on supervised and unsupervised learning tasks. Our hope is that adding noise to the network forces it to learning specific properties of the input distribution. These properties are crucial to the performance of supervised and unsupervised learning tasks.

Once the network has learnt interesting characteristics of the empirical distribution, our hope is that it has learnt to generalize these properties across input examples. An example of such a phenomenon can be found in [8], where a deep network created using stacked autoencoders learns properties of a distribution of images of digits. The network is able to fix digits like 6 and 7 of the input distribution, strongly suggesting that training using denoising autoencoders helped it generalize specific properties of the input digit image distribution. We argue that a similar phenomenon should be observed for gene expression profiles.

### Evaluating performance of deep networks using clustering

To measure the efficacy of the deep network in learning properties of the input distribution, we adopt the approach used by [15] and turn to the task of gene clustering. The task of gene clustering has important applications, like the following:

- It is likely for genes involved in the same cellular processes to be co-expressed. Since clustering favors grouping co-expressed genes in the same cluster, it becomes easier to study clusters of genes for their roles in processes.
- Genes in the same cluster are likely to be similar in coexpression.
- Genes clustered together are likely to be involved in a regulator pathway. This is because co-expression is a precursor to direct regulatory relationships.

We first cluster raw input using a fixed set of clustering algorithms. We then cluster samples regenerated from the deep network. Using pre-assigned cluster labels for all genes, the clustering performance of both sets of data is compared. We then report the performance of both sets of data on various network configurations. We present the details of the exact algorithm and clustering performance measure in the next section.

## Experiments

In this section, we describe our experimental setup and the details of datasets used. We also present a detailed explanation of the criteria used for assessing the performance of the deep architecture.

### Datasets

To test our model on real-world data, we used two datasets used by [15]. These datasets consist of time-series gene expression for two different sets of genes for yeast. They also include partition-clustering based labels for genes; each label instance corresponding to the cluster that gene belongs to.

Salient features of both datasets are described below:

- *Yeast cell cycle dataset 1*
- 
  A dataset capturing gene expression for the yeast cell cycle.
  Consists of 17 time ticks for a set of 384 genes made available in [15].
  External cluster labels partition the 384 genes into 5 clusters.
- *Yeast cell cycle dataset 2*
- 
  Consists of 17 time ticks for a set of 237 genes made available in [15].
  External cluster labels partition the 237 genes into 4 clusters.

### Evaluation criteria

To compare results of clustering against certain goldstandard labels, we need a measure of comparing two separate sets of cluster labels. Specifically, we need to qualitatively compare two separate instances of partition-based clustering on the same data set. We use the approach used in [15] and use the Adjusted Rand Index [18][19]. The adjusted rand index takes two sets of clusters, and checks whether every pair of items are in the same cluster in both the sets of clusters or not. This allows for it to compare two sets with different cardinalities as well. The expected value of the adjusted rand index between random sets of clusters is 0. It can take a maximum score of 1. The reader is advised to peruse [18] [19] for more details on the adjusted rand index.

A high adjusted rand index score means that the clustering result is in good agreement with the gold standard labeling. We compare the adjusted rand index scores of different datasets and report them in the following section.

### Clustering algorithms

We consider only partition-based clustering algorithms for evaluation purposes. Specifically, we use the k-means and the spectral clustering algorithm. Both algorithms require the number of clusters as input. Both algorithms were used via the SciPy library in Python.

### Hyperparameter tuning

Our experiments had a lot of parameters that required tuning. We considered different factors like batch size, number of training epochs, number of nodes in the hidden layer, corruption level and learning rate for SGD. Choosing the right set of parameters for a neural network is a difficult problem. A general rule of thumb is to not let the size of the hidden layer go below a certain threshold. This can be argued by considering the lossy compression that is achieved with a small hidden layer. Another rule of thumb used is to let the size of the hidden layer be equal to he number of principal components that capture the majority of the variance in the data. The most popular method to choose hidden layer is k-fold cross validation for a set of candidate hidden layer sizes. Care must be taken, however, to avoid overfitting to training data.

We noticed that the training error started settling after 1500 epochs, and chose a compatible number of training epochs. We kept the learning rate to be low in order to observe the convergence of the learning algorithm. We also kept the batch size to be a low number in order to take many gradient steps within a single epoch. Because denoising autoencoders are generally used for dimensionality reduction, we did not let the size of the hidden layer to be more than input layer size. For corruption level, we chose a range of values from no corruption to 20% corruption in the data. Since the datasets under consideration were not very large, we considered all possible combinations of the parameter values mentioned in table 1, and report the best results. We observed that most of the fluctuations in our results originated from changing the hidden layer size and corruption level.

Parameter setting for yeast-data experiments

TABLE I. PARAMETER SETTING FOR YEAST-DATA EXPERIMENTS
| Parameter settings |  |
| --- | --- |
| Parameter | Values used |
| Batch Size | 4,8,12 |
| Number of training epochs | 2000,5000 |
| Number of hidden nodes | 4,5,6,7,8,9,10,11,12,13,14,15,16,17 |
| Corruption level | 0,0.05,0.1,0.15,0.2 |
| Learning rate | 0.05,0.1 |

### Implementation

Code for all experiments was written in the Python library Theano [17]. All experiments were run on a machine with 2 Ghz of processing power and 16 GB of RAM. Since we were able to empirically observe improvements in performance with only one layer and training time for three layers was significantly higher, we used only one hidden layer.

Our argument is that using more layers, while taking phenomena like overfitting into account, will only improve performance further, but a significant cost of computation power and time. For all optimizations, we used symbolic differentiation of Theano for gradient calculations and used Stochastic Gradient Descent (SGD) to optimize the crossentropy objective function.

## Results

In this section, we discuss the results of experiments on both Yeast datasets together. We consider various aspects of the denoising autoencoder and their effects on clustering performance. We also consider the performance of PCA versus our methods.

### Raw data vs regenerated data

The results of experiments on both the yeast datasets are presented in figs. 2 and 3. In both cases, it is clear that regenerated data outperforms raw data. This can possibly be explained by the unique training procedure of the architecture, which involves corrupting data partially in order to force the network to learn important relationships in the underlying distribution.

**Fig. 2.**
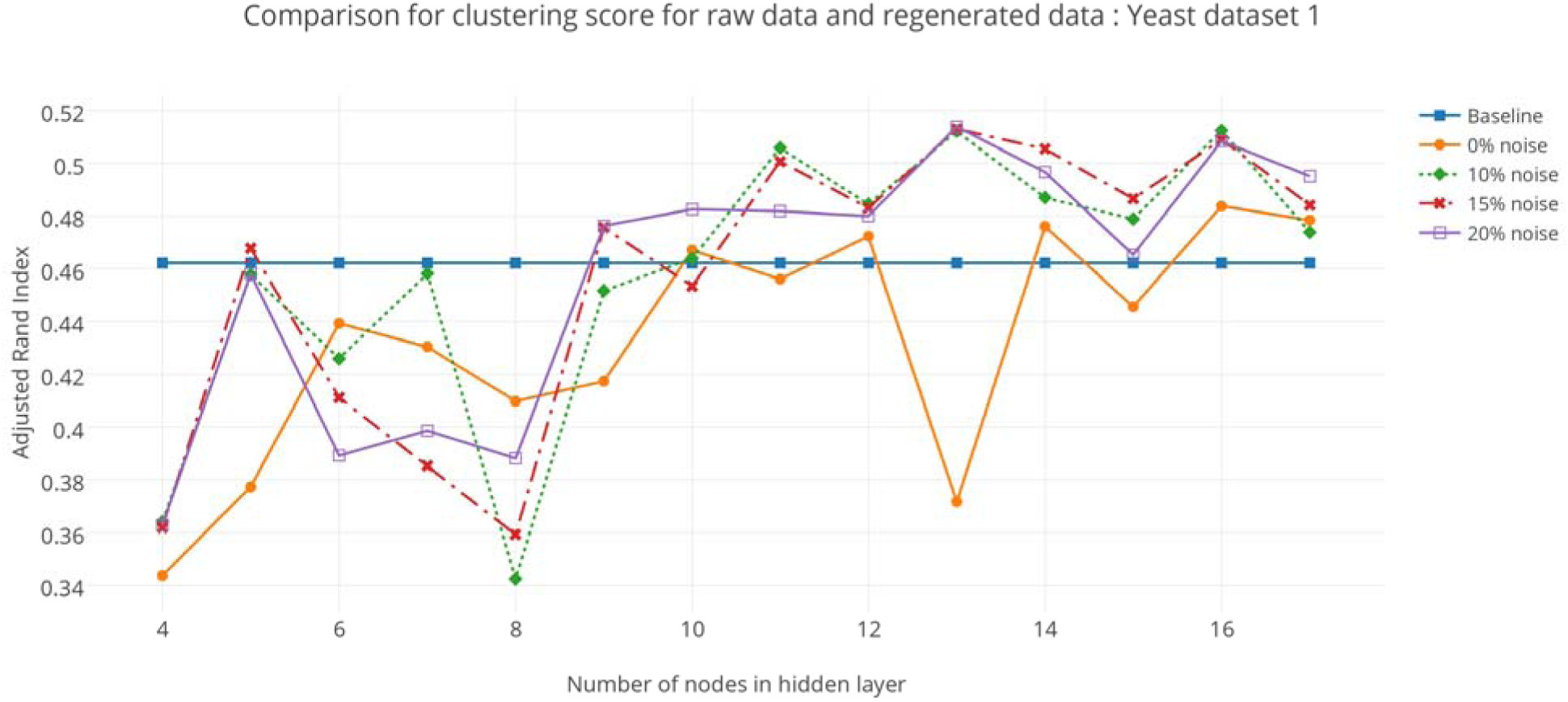
Performance comparison of raw data of yeast dataset 1 and samples regenerated from the trained network on the task of clustering.

**Fig. 3.**
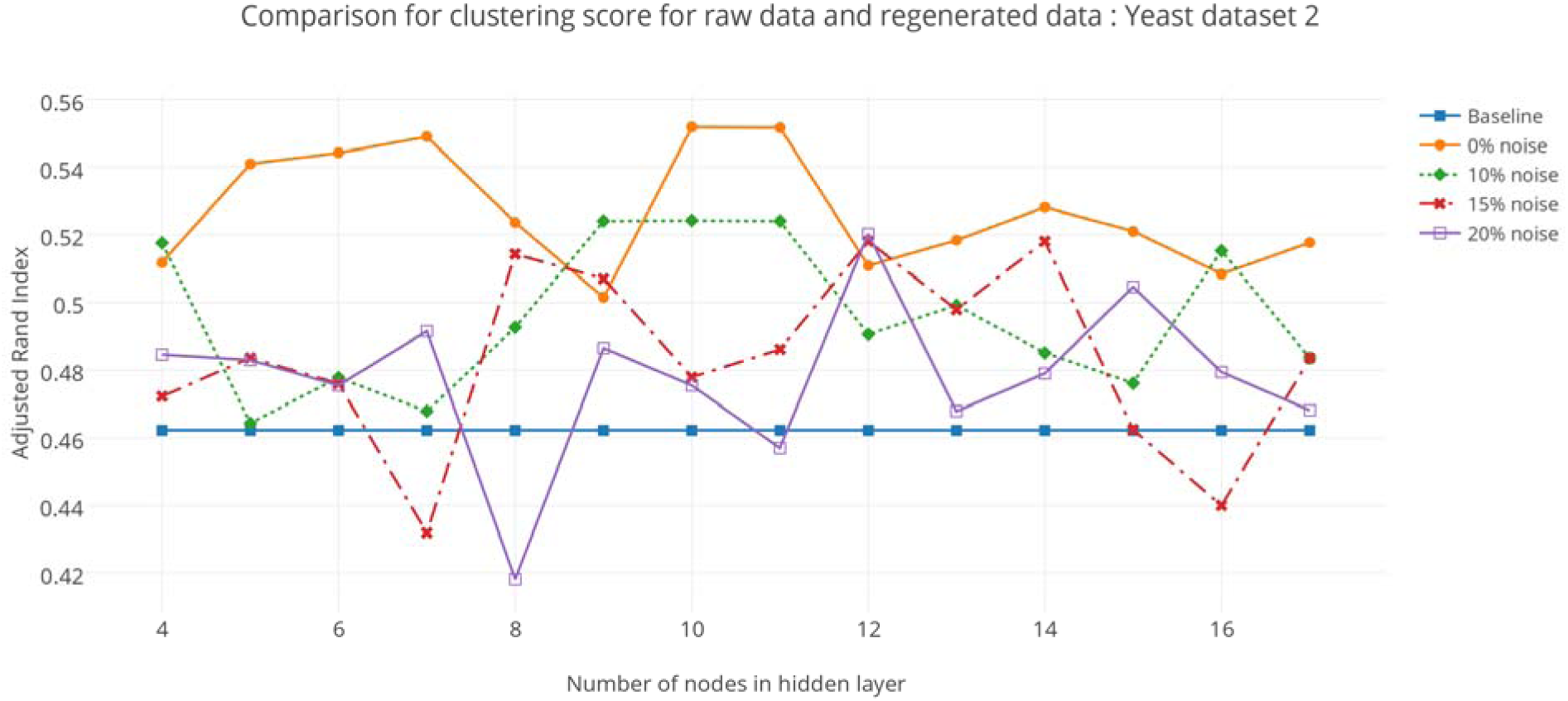
Performance comparison of raw data of yeast dataset 2 and samples regenerated from the trained network on the task of clustering.

### PCA vs regenerated data vs raw data

The results of clustering with principal components are presented in figure 4. We chose to compare the first k components, and also the performance of randomly chosen components, with the performance over raw input data. The performance with principal components is quite poor compared to both raw and regenerated data. We conclude that principal components do not necessarily capture relationships among the components of the input, and are not suitable to learn the intricacies of the underlying distribution.

**Fig. 4.**
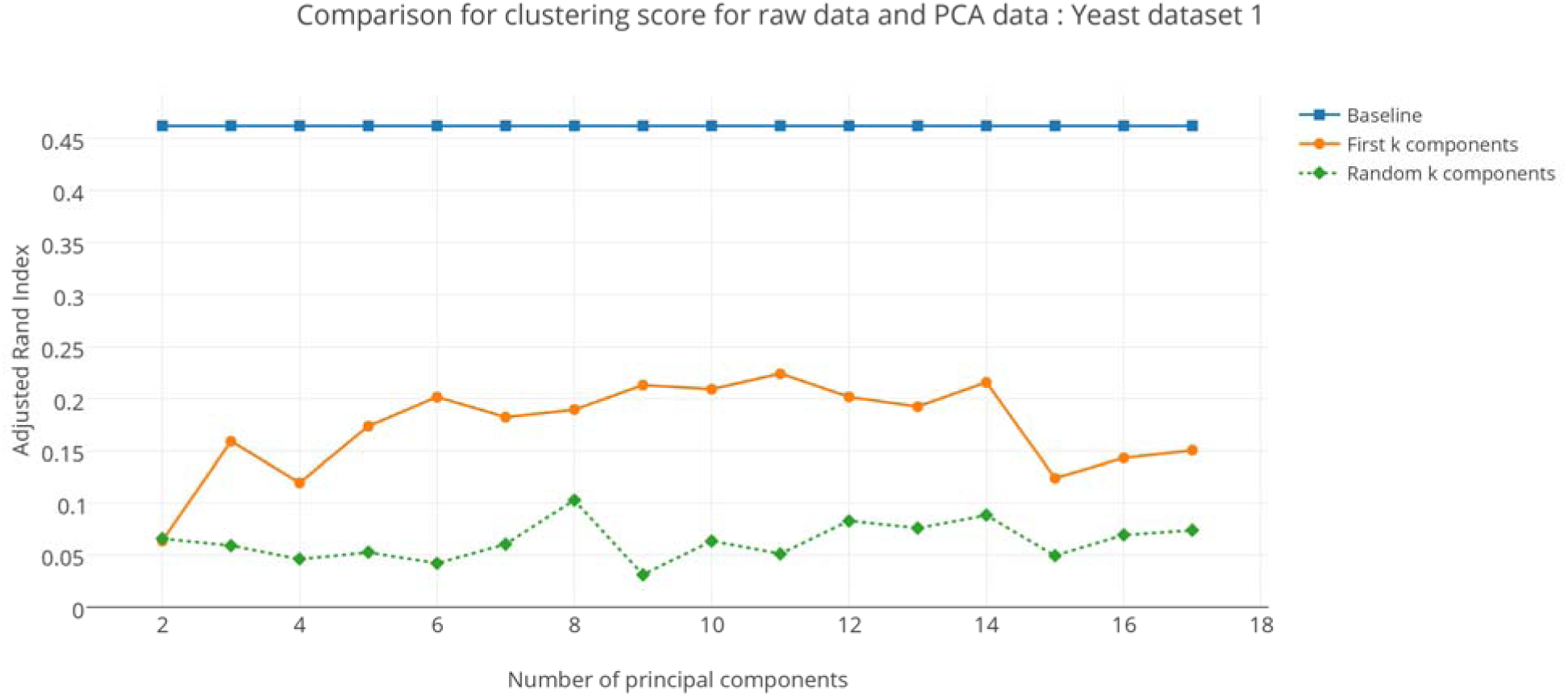
Performance comparison of raw data of yeast dataset 1 and principal components of data. The results compare performance of raw data versus the first k components, or a random number of components from all principal components.

### Effect of percentage of training noise

In both Fig. 2 and Fig. 3, it is clear that clustering performance improves when at least 5 % noise is used. The performance on using 0% noise, when the input is not corrupted at all during training, is much less clear - it improves in Fig. 3 and does worse than raw data in Fig. 2. However, performance with any degree of noise is better than the baseline performance of raw data, provided that enough number of hidden nodes have been used. The distinction between performances of different noise levels is not very clear.

### Effect of number of nodes in hidden layer

The results of fig.2 make it clear that clustering performance improves as the number of nodes in the bottleneck layer is improved. A possible reason for this to happen could be the network’s ability to retain more information when the number of nodes in the low-dimensional code is increased. A similar trend is also observed for the second dataset in Fig. 3, although the trend is not as strongly evident.

## Conclusions and Future Work

Our experiments demonstrate the empirical effectiveness of using deep networks as a pre-processing step for clustering of gene expression data. Deep networks are initialized using denoising autoencoders to learn interesting properties of gene expression profiles. Clustering of genes using gene expression data is an important task for research related to interactions and regulation among genes. Thus, we empirically demonstrate the advantage of using gene expression samples regenerated from the low-dimensional codes for the task of clustering. We argue that this process works because denoising autoencoders do not merely perform the task of retaining information about the distribution, but also generalize important and interesting properties of the input distribution across all input samples. This is made possible by the unique training strategy of denoising autoencoders.

In our experiments, we demonstrate the efficacy of using regenerated samples for the task of clustering genes into groups. Our empirical results indicate that in general, even a shallow network can outperform both raw data and PCA on the task of partition-based clustering. This was observed consistently across both datasets under consideration. In the case of the first dataset, we observe that adding noise during training significantly boosts system performance. On the other hand, adding training noise to the second dataset does improve system performance. However, the distinction between the performances of different noise levels is much less clear. Since we constrain the size of the hidden layer to be small, even training without noise may occasionally help the network learn interesting properties of the distribution. on the whole, regenerated samples outscored raw data on the task of partition-based clustering significantly.

In this work, we have demonstrated only one application of regenerating samples from deep networks. We also use a fairly simple architecture. In future work, we aim at using more datasets and deeper architectures, pre-trained greedily followed by global optimization. We also aim at using the lowdimensional coding learned by the deep network as features for supervised learning tasks like protein-protein interaction prediction.

## Acknowledgment

AG is grateful to Volkan Cirik for valuable discussions on the capabilities of autoencoders. This work has been funded by the Biobehavioral Research Awards for Innovative New Scientists (BRAINS) grant R01MH094564 awarded to MKG by the National Institute of Mental Health of National Institutes of Health (NIHM/HIM) of USA.

